# The genetic basis for PRC1 complex diversity emerged early in animal evolution

**DOI:** 10.1101/2020.03.18.997064

**Authors:** James M Gahan, Fabian Rentzsch, Christine E Schnitzler

## Abstract

Polycomb group proteins are essential regulators of developmental processes across animals. Despite their importance, studies on Polycomb are often restricted to classical model systems and, as such, little is known about the evolution of these important chromatin regulators. Here we focus on Polycomb Repressive Complex 1 (PRC1) and trace the evolution of core components of canonical and non-canonical PRC1 complexes in animals. Previous work suggested that a major expansion in the number of PRC1 complexes occurred in the vertebrate lineage. Here we show that the expansion of the PCGF protein family, an essential step for the establishment of the large diversity of PRC1 complexes found in vertebrates, predates the bilaterian-cnidarian ancestor. This means that the genetic repertoire necessary to form all major vertebrate PRC1 complexes emerged early in animal evolution, over 550 million years ago. We further show that *PCGF5*, a gene conserved in cnidarians and vertebrates but lost in all other studied groups, is expressed in the nervous system in the sea anemone *Nematostella vectensis*, similar to its mammalian counterpart. Together this work provides an evolutionary framework to understand PRC1 complex diversity and evolution and establishes *Nematostella* as a promising model system in which this can be further explored.

## Introduction

Polycomb group proteins were first described in *Drosophila* as genes essential for patterning during embryogenesis and were subsequently shown to play crucial roles in cell differentiation and the maintenance of cell fate during development in many systems (1). Polycomb group proteins establish “facultative” heterochromatin and are required to maintain repression of key developmental genes such as *Hox* genes. As such, loss of Polycomb often results in homeotic transformations due to misexpression of *Hox* genes (1). In addition, Polycomb proteins can maintain genes in a poised state, which is characterized by the simultaneous presence of distinct histone modifications that are associated with transcriptional repression and activation (2).This poised state allows the rapid activation of transcriptional programs and accordingly, the Polycomb system is not only required for repression but also for the temporal control of transcriptional activation during development (2). In addition to this, Polycomb proteins are frequently found mutated in cancer and represent a popular therapeutic target (3).

Polycomb proteins belong to one of two complexes: Polycomb Repressive Complex 1 or 2 (PRC1 or PRC2, respectively). PRC2 complexes catalyze trimethylation of lysine 27 on histone H3 (H3K27me3), a repressive histone modification (4). PRC1, on the other hand, ubiquitinates histone H2A and mediates chromatin compaction and gene silencing (5–13). The classical model of transcriptional silencing by Polycomb complexes entails first recruitment of PRC2, which deposits H3K27me3, followed by PRC1 recruitment through its H3K27me3 binding subunit, leading to H2A ubiquitination and repression (14–16). In recent years, this model has been elaborated upon extensively, revealing a more complex interplay between PRC1 and PRC2 components, histone modifications and other factors such as DNA methylation and CpG content that regulate the recruitment and activity of both complexes and subsequent transcriptional repression (17–31).

Both PRC1 and PRC2 are large, multi-subunit protein complexes. In *Drosophila*, PRC2 consists of a core of three proteins: Extra sex combs (Esc), Suppressor of Zeste 12 (Su(z)12) and Enhancer of Zeste (E(z)) (4) (see Table S1 for nomenclature of Polycomb proteins). PRC1 consists of four proteins: Sex combs extra (Sce or dRING), Posterior sex combs (Psc or its holomolg Su(z)2), Polyhomeotic (Ph) and Polycomb (Pc). Vertebrate PRC2 is highly similar to that of *Drosophila*, with EED, SUZ12 and EZH1/2 as the orthologs of Esc, Su(z)12 and E(z), respectively (4). PRC1, in contrast, is thought to have undergone an expansion in vertebrates, represented by a collection of related complexes each sharing a core consisting of RING1A or RING1B, vertebrate homologs of dRING, and one of the six vertebrate Polycomb Group RING Finger (PCGF) proteins, the homologs of *Drosophila* Psc (32). cPRC1.2 and cPRC1.4, the canonical PRC1 complexes, consist of either PCGF2 or PCGF4, respectively, in a complex with RING1A/B, one Chromobox protein (CBX, the homologs of *Drosophila* Pc) and one Polyhomeotic-like protein (PHC) (32) (Fig.1A). Further diversification within the vertebrate canonical complexes occurs due to the presence of five different potential CBX subunits (33–36), and three different PHC proteins (32). The non-canonical or variant PRC1 complexes, ncPRC1.1-1.6, consist of one PCGF protein, as well as RING1A/B, RYBP or its homolog YAF2 and other complex specific subunits (22, 32, 37–39) (Fig. 1A). The integration of either a CBX protein (in cPRC1) or RYBP/YAF2 (in ncPRC1) is based on their mutually exclusive interaction with RING1A/B (32, 33, 40, 41). The majority of H2A ubiquitination is mediated by the non-canonical complexes (42) while only the canonical complexes can be recruited by H3K27me3 through their CBX subunit (15, 16, 43) and have the ability to mediate both local compaction and long range interactions (6, 10–12, 44, 45). While complete loss of PRC1 via deletion of RING1A/B is lethal (46, 47), different PRC1 complexes can have distinct roles, owing to both the different subunits but also tissue specific expression of complex members (19, 22, 39, 48–56). In *Drosophila*, in addition to the canonical complex outlined above, two non-canonical PRC1 complexes have been described: dRAF, which contains KDM2, a lysine demethylase subunit (57), and a complex which contains an alternative Psc homolog (58).

**Fig. 1.**
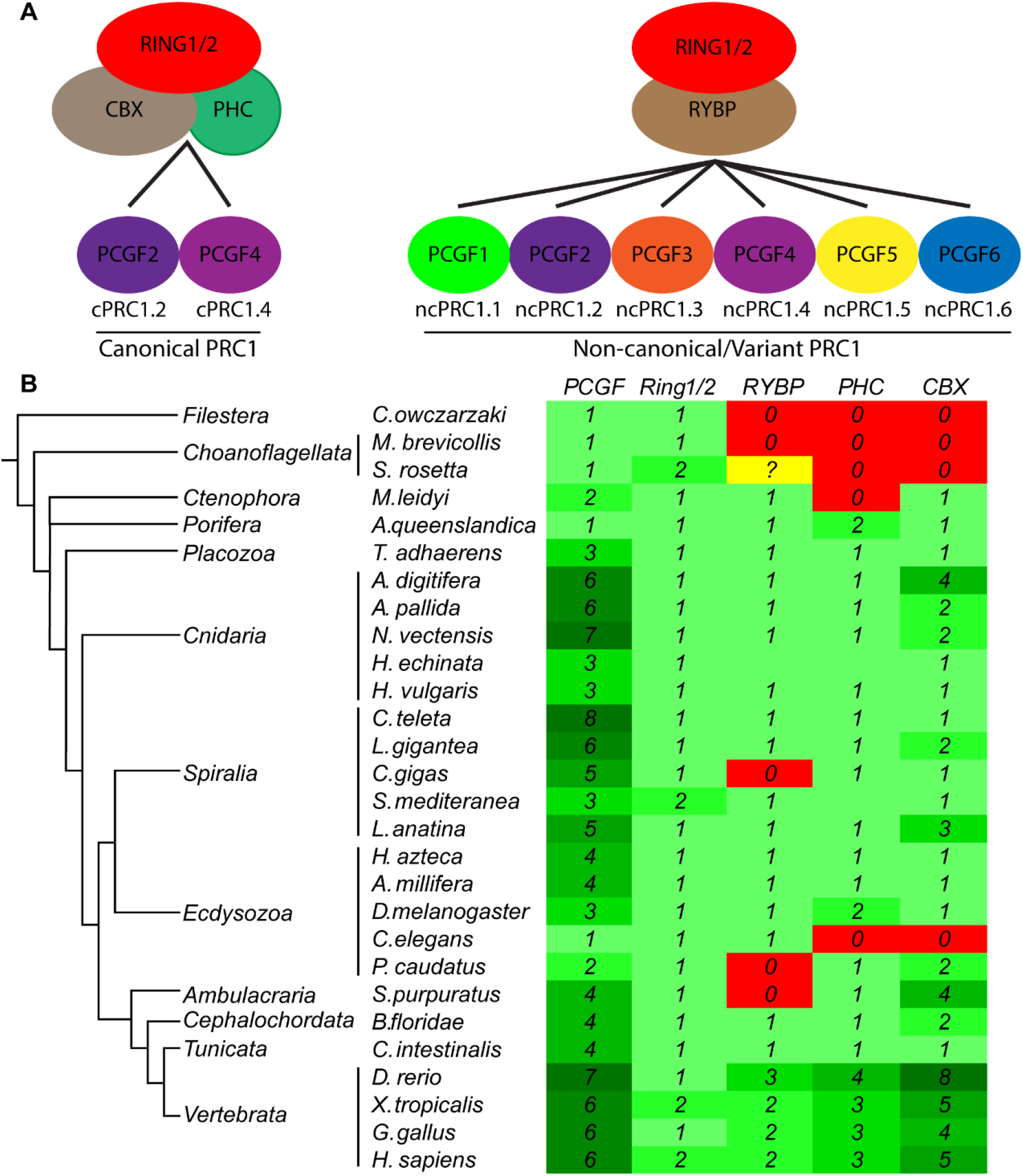
An animal-specific set of core PRC1 components. (A) Schematic showing the core components of all known PRC1 variants in vertebrates. (B) Table showing the presence or absence as well as the number of PCGF, CBX, PHC, RING1/2 and RYBP homologs in representative animal and single cell eukaryote species. Green indicates the presence of a homolog, red indicates absence and yellow indicates cases where there is ambiguity. The number of homologs in a particular species is indicated both by number but also by intensity of green colour. The full set of the PRC1 components is only found in animals.

PRC1 complexes containing RING1/2 and PCGF proteins are present in plants, but many of the other components in these complexes are distinct to those in animals (59–62). Similarly, RING1/2 and PCGF are encoded in the genomes of many unicellular eukaryotes like choanoflagellates, ichthyosporeans, and filasterians (1), but it is not known whether they form complexes with PRC1-like functions. Central to the current understanding of the evolution of PRC1 complexes, previous analyses have shown that compared with *Drosophila*, vertebrates have an expanded number of CBX, PHC, and PCGF proteins. This supported a scenario in which the diversity of PRC1 complexes mainly arose in vertebrates (63, 64).

Here we searched a broad selection of animal and closely related unicellular eukaryote genomes for the presence of genes encoding the core proteins required to make all possible PRC1 complexes described above and performed a phylogenetic analysis on PCGF proteins to understand their evolution. While we find the expansion of CBX and PHC proteins in vertebrates is likely correct, we determined that, contrary to current thinking, the diversity found in mammalian PCGF proteins emerged more than 550 million years ago, before the last common ancestor of bilaterians and cnidarians (65). Thus, the genetic basis for PRC1 complex diversity appeared early in animal evolution but has been lost secondarily in different animal lineages. Using a transgenic reporter line in the anthozoan cnidarian *Nematostella vectensis*, we further show that *PCGF5* genes may have ancient roles in the nervous system.

## Results

We searched 28 genomes, representing diverse animal clades and the two closest unicellular outgroups to animals, choanoflagellates and filasterians, for the presence of homologs of the core components of canonical or non-canonical PRC1, i.e. *RING1/2* (genes encoding RING1A/B), *PCGF*, *CBX*, *PHC*, and *RYBP*, using either *Drosophila* or human sequences as query (see Materials and Methods). The presence/absence as well as the number of genes per species are shown in Figure 1B (see also (1)). There are single copies of *RING1/2* in most species with the exception of some vertebrates where there are two copies, *RING1A* and *RING1B*, and in the platyhelminth *Schmidtea mediterranea* where there are also two *RING1/2* genes. We found no CBX and PHC genes outside animals and both genes were lost in the lineage leading to the nematode *C. elegans.* We identified only one copy of PHC in most animals except vertebrates where we find three copies, the sponge *Amphimedon queenslandica* which has two genes, and *Drosophila melanogaster* which has two almost identical PHC genes, the result of a recent duplication event (66). For the *CBX* genes, we found a relatively large diversity in gene number (ranging from one to eight) in different animals. *RYBP*, in contrast, is present as a single copy gene in most invertebrates, but is represented by two paralogs in vertebrates, named *RYBP* and *YAF2*. Some invertebrate species (the oyster *Crassostrea gigas*, the priapulid worm *Priapulus caudatus*, and the sea urchin *Strongylocentrotus purpuratus*) lack an *RYBP* homolog, likely due to secondary loss as these species are only distantly related to each other. Interestingly, a putative homolog of *RYBP*, the unique component of non-canonical PRC1, can be found in the choanoflagellate *Salpingoeca rosetta*, but not in another choanoflagellate, *Monosiga brevicollis*, and also not in the filasterean *Capsaspora owczarzaki* as previously noted (1). The level of sequence similarity of the *S. rosetta* gene compared to animal RYBP genes is, however, very low and it does not contain the Yaf2/RYBP C-terminal binding motif which is present in all other *RYBP* genes. While it is possible that there was a *RYBP* gene present in the last common ancestor of choanoflagellates and animals that did not contain a Yaf2/RYBP C-terminal binding motif, we prefer to label this *S. rosetta* gene as a putative RYBP gene (shown in (Fig. 1B) as a question mark).

Surprisingly, we found a wide range in the total number of *PCGF* genes per animal (Fig. 1B). Previous work had shown that the *PCGF* family expanded only in vertebrates but we found 6-7 *PCGF* genes in anthozoan cnidarians and eight in the annelid *Capitella teleta*, more than found in mammals. This diversity in the number of *PCGF* genes in each animal genome we searched suggests many lineage specific gains and/or losses.

Thus, in contrast to RING1/2, PHC, and RYBP genes, the number of *PCGF* genes varies considerably among animals. This observation prompted us to use phylogenetic analyses to understand the evolution of the PCGF gene family in more detail. We performed a phylogenetic analysis on the full set of taxa in (Fig. 1B) using PCGF and RING1/2 proteins as an outgroup or PCGF proteins alone using both maximum likelihood and Bayesian methods (Fig. S2-S5). We also ran the analysis on the PCGF and RING1/2 proteins with a reduced set of sequences corresponding to cnidarian and selected bilaterian lineages (Fig. 2) and Fig. S1). In all cases, the overall topology of the tree was similar. We found that the *PCGF* genes fall into five families, which we termed PCGF1, PCGF2/4, PCGF3, PCGF5, and PCGF6 based on the vertebrate homologs present in the groups. The “canonical” *PCGF2/4* and the “non-canonical” *PCGF1, 3, 5, 6* genes form sister groups, with additional subgrouping of the “non-canonical” genes into PCGF1, PCGF3, PCGF5, and PCGF6 subgroups. Figure 3A summarizes the presence of genes within the different families in all bilaterian and cnidarian species studied with the exception of *Ciona intestinalis* as we could not confidently assign some genes from this species. All of the *PCGF* families contain sequences from bilaterian and cnidarian genomes indicating they originated before the split of these two major animal groups. All but the *PCGF5* group also contain sequences from both protostomes (Ecdysozoa and Spiralia, see Fig. 1) and non-vertebrate deuterostomes. Although many species have more than one gene within the *PCGF2/4* clade, it is likely that these arose through lineage specific duplications. This is the case for vertebrate *PCGF2* and *PCGF4* (*Bmi1*) as well as for *Drosophila PSc* and *Suz(2)* and the two *PCGF2/4* genes present in anthozoan cnidarians. There have also been extensive losses of many *PCGF* genes, most strikingly that of *PCGF5*, which has been lost at the base of the protostomes but also in the non-vertebrate deuterostomes studied here. Additional losses have occurred in specific lineages, for example loss of *PCGF6* in Ecdysozoa and *PCGF3* in hydrozoan cnidarians (Fig. 3A). Among the analyzed taxa, anthozoan cnidarians (three species) and vertebrates (four species) are the only ones in which all five PCGF subgroups are present.

**Fig. 2.**
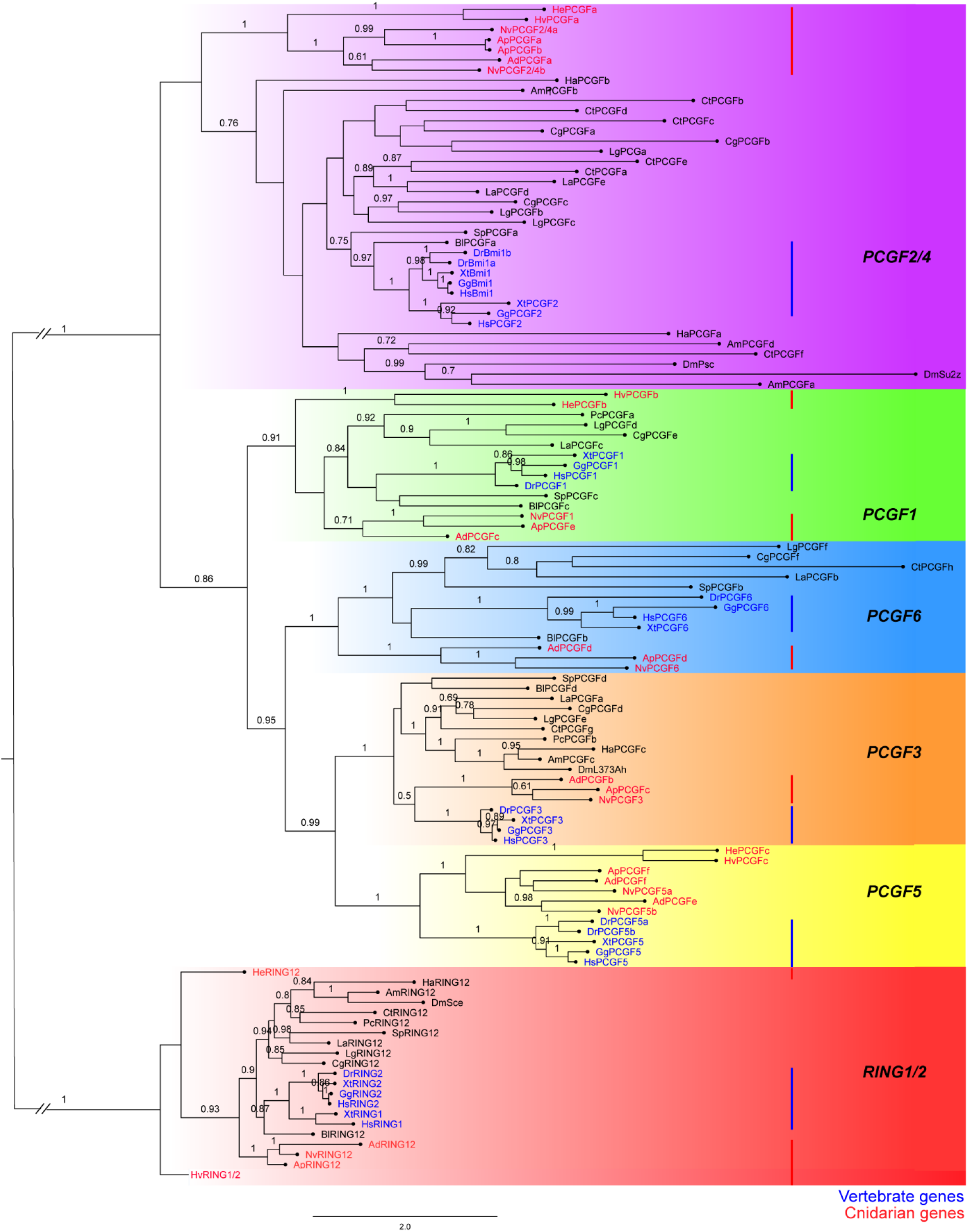
Subgroups of *PCGF* genes emerged early in animal evolution. Phylogenetic analysis of cnidarian and bilaterian *PCGF* and *RING1/2* genes according to Bayesian analysis using *RING1/2* genes as outgroup. There are five major families of *PCGF* genes (*PCGF1, 2/4, 3, 5, 6*), which are highlighted by different colored boxes. Numbers above branches correspond to Bayesian posterior probabilities. Only values ≥ 0.7 are shown. Red bars and red font indicate the position of vertebrate genes and blue bars and blue font indicates the position of cnidarian genes. Species names are abbreviated as follows: Ad, *Acropora digitifera*; Ap, *Aiptasia pallida*; Am, *Apis mellifera*; Bl, *Branchiostoma floridae*; Cg, *Crassostrea gigas*; Ct, *Capitella teleta*; Dr, *Danio rerio*; Dm, *Drosophila melanogaster*; Gg, *Gallus gallus*; Ha, *Hyalella azteca*; Hs, *Homo Sapiens*; He, *Hydractinia echinata*; Hv, *Hydra vulgaris*; La, *Lingula anatina*; Lg, *Lottia gigantea*; Nv, *Nematostella vectensis*; Pc, *Priapulus caudatus*; Sm, *Schmidtea mediterranea*; Sp, *Strongylocentrotus purpuratus*; Xt, *Xenopus tropicalis.*

**Fig. 3.**
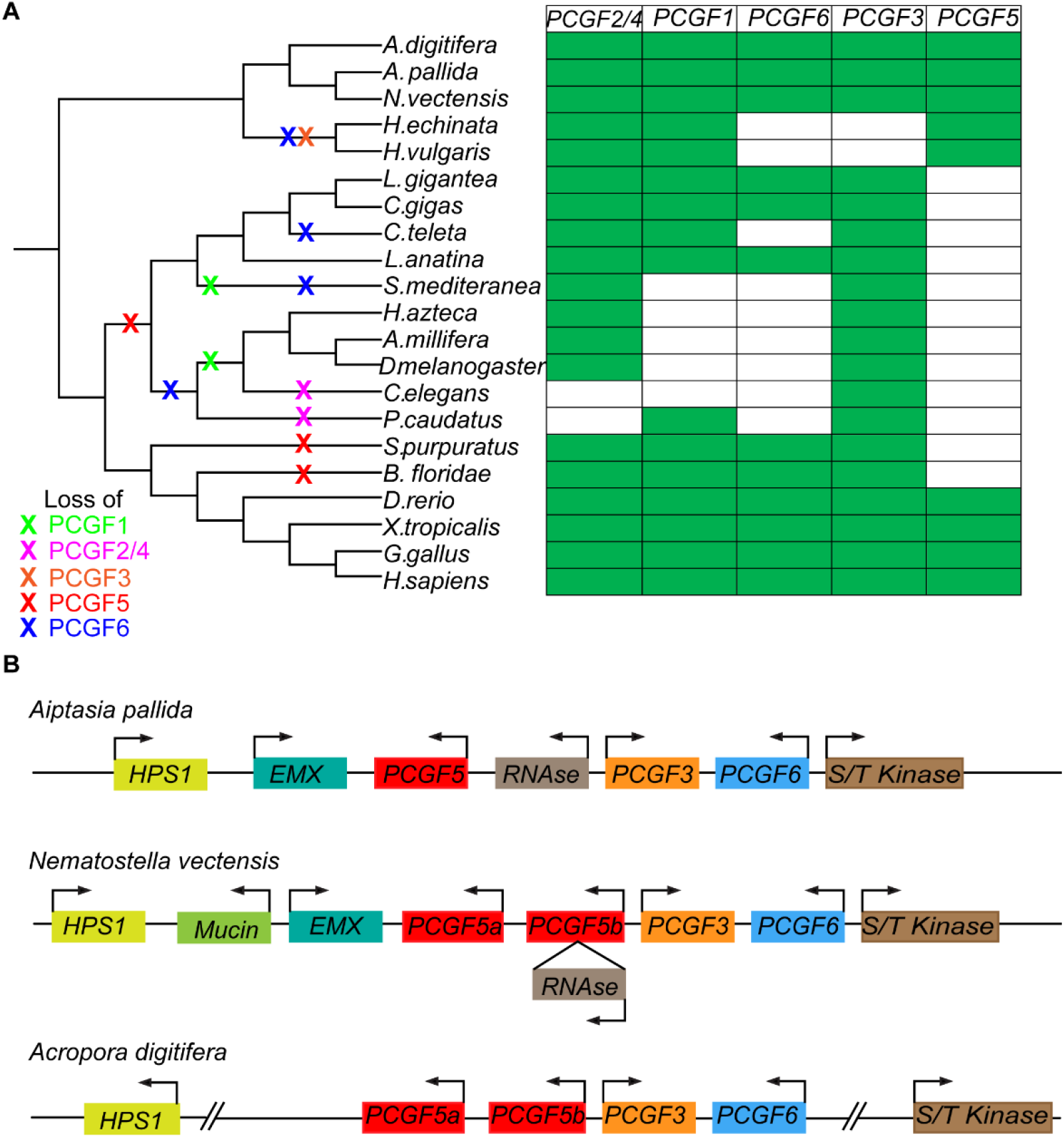
Anthozoan cnidarians have a complete complement of PCGF subgroups and a cluster of “non-canonical” PCGFs. (A) Table showing the presence or absence of members of the various PCGF families identified in the genomes of all cnidarian and bilaterian species studied here except *Ciona intestinalis* which was excluded due to the fact we could not clearly place all its PCGF homologs unambiguously into one of these families. An X on the tree indicates predicted gene losses. (B) Schematic representation depicting the *PCGF* gene cluster and gene synteny between *Nematostella vectensis*, *Aiptasia pallida*, and *Acropora digitifera*. The PCGF genes were named based on their position in the phylogeny (Fig. 2). Other genes in the cluster were named based on the closest BLASTp hit in human. HPS1: Hermansky-Pudlak syndrome 1 protein, Mucin-22: Mucin-22 isoform-1 precursor, EMX: homeobox protein EMX1, S/T kinase: Leucine rich repeat serine/threonine-protein kinase 2, RNase: probable ribonuclease ZC3H12B.

The position of the genes from the unicellular groups (choanoflagellates and filasterians) as well as other non-bilaterian animal groups (ctenophores, sponges, placozoans) was ambiguous on the trees. The two choanoflagellate *PCGF* genes fall either within the PCGF5 clade or as sister to the *PCGF5* clade depending on whether RING1/2 genes are included in the analysis (Fig. S2-S5). This may be due to some ancestral characteristics of PCGF being retained in the *PCGF5* genes or alternatively due to convergence. Similarly, the single *PCGF* gene in the filasterian *Capsaspora* has a shifting position within the trees (Fig. S2-S5). In neither case do these positions have a high level of support. A similar situation is seen for some or all of the genes from the ctenophore *Mnemiopsis leidyi*, the placozoan *Trichoplax adhaerens*, and the sponge *Amphimedon queenslandica* (Fig. S2-S5). Thus, it is not possible from this analysis to confidently derive conclusions about *PCGF* gene evolution before the last common cnidarian-bilaterian ancestor.

Anthozoan cnidarians, an animal clade containing corals and sea anemones, are the earliest-diverging animals that have at least one member of each of the PCGF families and indeed are the only group outside vertebrates to have this. We therefore sought to investigate this group further. We particularly focused on *Nematostella vectensis*, the starlet sea anemone, due to the availability of experimental tools (67). While analyzing the *PCGF* complement in *Nematostella* we noted that four of the *Nematostella* genes are arranged in a genomic cluster: *NvPCGF5a*, *NvPCGF5b*, *NvPCGF3* and *NvPCGF6* (Fig. 3B). We then looked in other anthozoan genomes and found the genomic cluster to be conserved in both *Aiptasia*, another sea anemone, and *Acropora*, a coral (Fig. 3B). We found no evidence for conservation of this cluster in vertebrate or amphioxus genomes. Interestingly, the order of the genes along the cluster in anthozoans reflects their evolutionary relationships that we found in our trees: the *PCGF5* genes (there are two paralogs in *Nematostella* and *Acropora*) are located next to each other, the most closely related *PCGF3* is located adjacent to the *PCGF5* genes and the more distantly related *PCGF6* is located on the other side of *PCGF3*.

To investigate whether distinct *PCGF* genes in *Nematostella* may have distinct functions we sought to analyze their expression. We first interrogated a previously published developmental time course (68), which integrates RNAseq data from several studies (69–71). We saw that both “canonical” *PCGF* genes, *NvPCGF2/4a* and *NvPCGF2/4b*, had similar expression dynamics during development although with different levels (Fig. S6). In the case of the “non-canonical” *PCGF* genes, we found that there is substantial variability in their expression (Fig. 4A). *NvPCGF5a*, for example, is not maternally expressed and its expression reaches maximum levels at around planula larva stage (approximately 48hrs post fertilization) while *NvPCGF5b* is maternally expressed and reaches the same level at larva stage as *NvPCGF5a* but with higher expression during early embryonic stages. *NvPCGF3* is also maternally expressed and its levels remain steady until blastula stages when its levels drastically increase before plateauing. Both *NvPCGF1* and *NvPCGF6* are highly expressed maternally and high levels of both genes are maintained during early embryogenesis before levelling out at a lower level after gastrulation.

**Fig. 4.**
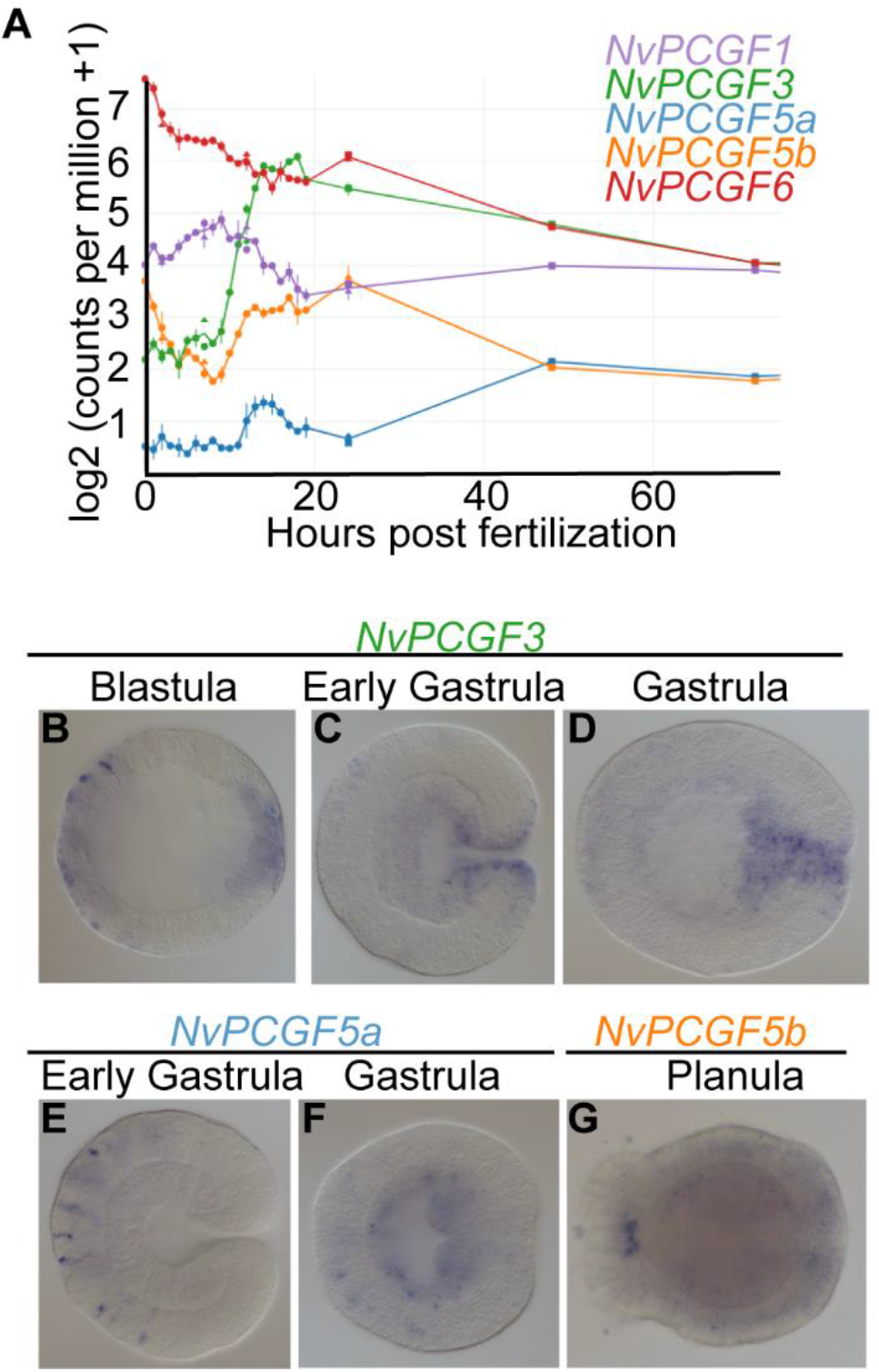
Temporally and spatially dynamic expression of non-canonical *PCGF* genes in *Nematostella vectensis*. (A) Expression analysis of the *Nematostella* non-canonical *PCGF* genes throughout embryonic development taken from NvERTx database (68). (B-G) RNA in situ hybridization of *NvPCGF3* (B-D), *NvPCGF5a* (E) and *NvPCGF5b* (F-G) at indicated developmental time points. Lateral views with aboral pole to the left.

Some vertebrate *PCGF* genes display spatial expression patterns, with higher levels in specific tissues or cells (72–74). We performed RNA in-situ hybridization at different developmental stages to determine whether such spatial regulation also occurs for *Nematostella PCGF* genes. *NvPCGF3* is expressed at blastula stage in two distinct domains on opposing sides of the embryo (Fig. 4B), presumably corresponding to the oral and aboral poles. At gastrula stage, this pattern continues with expression being localized to oral and pharyngeal tissue and, less pronounced, to the aboral pole (Fig. 4C, D). *NvPCGF5a* can first be detected by in-situ hybridization at early gastrula stage when it is expressed in scattered cells on the aboral side of the embryo (Fig. 4E). This expression pattern continues into later stages, although weaker, and spreads into the endoderm (Fig. 4F). Localized expression of *NvPCGF5b* is first detectable at planula stage when it is expressed in the apical tuft, albeit very weakly (Fig. 4G). We note that the RNAseq data (Fig. 4A) show that both *NvPCGF5* paralogs are also expressed at stages at which we cannot detect them by in-situ hybridization, potentially due to low level and/or broad expression at those stages. We were unable to find distinct/localized expression patterns for the other PCGF genes.

Given that the *NvPCGF5a* expression pattern is similar to that seen for neural genes at these stages (75–77) and that vertebrate *PCGF5* is highly expressed in neural progenitors (56, 72) we wanted to investigate further these *NvPCGF5a* expressing cells. To do this we generated a transgenic reporter line expressing *eGFP* under the control of the *NvPCGF5a* regulatory elements. *eGFP* could be detected in these animals in scattered cells in the aboral half of the embryo from gastrula stage on (Fig. 5A-C) and later additionally at lower levels throughout the aboral tissue (Fig. 5B-C). The morphology of the scattered cells matched that expected of neurons and/or sensory cells with many cells seen with an apical cilium and basally branching neurites. We went on to cross this line to other published neuronal reporter lines. This revealed that the *NvPCGF5:*:*eGFP*^+^ cells represent a subpopulation of both the *NvFoxQ2d*::*mOrange*^+^ positive sensory cells (78) (Fig. 5D) and the *NvElav1*::*mOrange*^+^ positive neurons (79) (Fig. 5E). Together these data show that *NvPCGF5a* is expressed in a subset of neural cells in *Nematostella.*

**Fig. 5.**
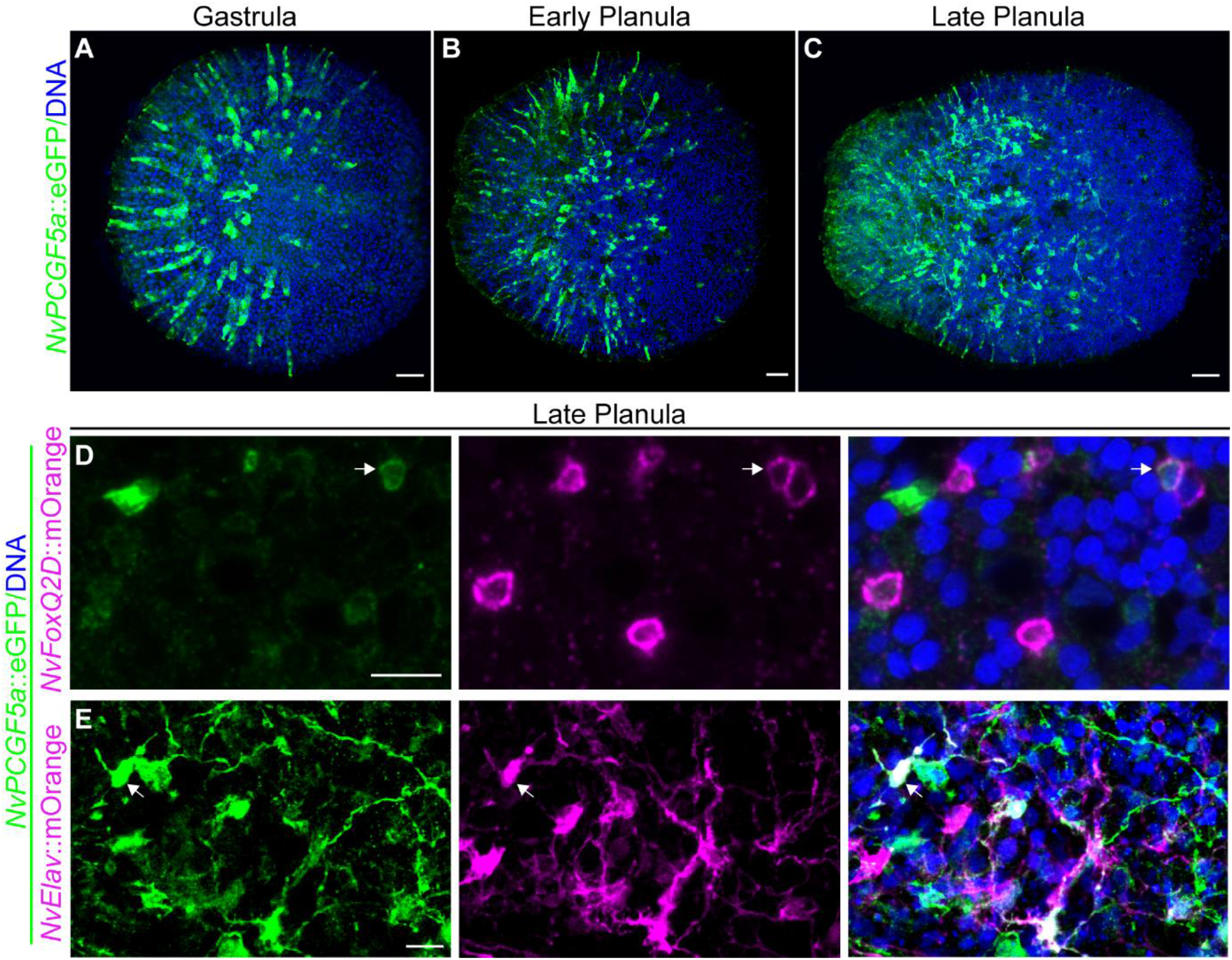
An *NvPCGF5a*::eGFP reporter line marks a subset of neural cells in *Nematostella*. (A-C) Immunostaining for eGFP highlights neural cells in the aboral part of *Nematostella* embryos at gastrula (A), early planula (B) and late planula. Lateral views with aboral pole to the left. (D, E) Immunostaining on late planula of double transgenic animals bearing *NvPCGF5a*::eGFP and *NvFoxQ2d*::mOrange (D) or *NvElav:*:mOrange(E) transgenes shows co-localization in a subset of cells in both cases. Scale bar: 20 μM (A-C), 10 μM (D, E).

## Discussion

Here we analyze the evolution of the core components of canonical and variant PRC1 in animals. PRC1 complexes were thought to have experienced a diversification in vertebrates, mainly due to expanded repertoires of CBX and PCGF genes (56, 61). We show that although some expansion of PRC1 components did indeed occur in vertebrates, i.e. expansion of *CBX* and *PHC* genes, the expansion of the *PCGF* gene family occurred much earlier, before the last common ancestor of cnidarians and bilaterians. Our analysis indicates that there were likely five PCGF proteins in the last common ancestor of bilaterians and cnidarians and that there was only one subsequent duplication, within the *PCGF2*/4 family, in the lineage leading to vertebrates. This is an intriguing finding as PCGF proteins define the composition and identity of the main canonical and non-canonical PRC1 complexes (32). We show that the non-canonical *PCGF* genes (those encoding PCGF1, 3, 5 and 6) are more closely related to each other than to the canonical *PCGF2*/*4* family and that at least PCGF3, 5 and 6 arose from sequential duplications of an ancestral gene. This is evident in anthozoan cnidarians where *NvPCGF5*, *NvPCGF3* and *NvPCGF6* genes have been maintained in a genomic cluster. While the presence of this cluster in anthozoans is informative for the evolution of *PCGF* genes, it remains to be determined which genomic features favored its retention in this particular clade of animals.

The previously assumed expansion of Polycomb complexes in vertebrates has been deduced primarily from comparisons to *Drosophila* and *C. elegans*. *Drosophila* has only three *PCGF* genes, two belonging to the PCGF2/4 family and one to the PCGF3 family. Despite the lack of a *PCGF1* homolog, there is a PRC1 complex in *Drosophila*, dRAF, which resembles vertebrate ncPRC1.1 in that it contains the lysine demethylase KDM2, but differs from ncPRC1.1 by the presence of a PCGF2/4, rather than a PCGF1 protein (57). This may suggest that non-canonical PRC1 complexes can switch PCGF components over evolutionary time or it may be a specific case caused by the loss of PCGF diversity in the lineage leading to *Drosophila*. A recent report has found that the *Drosophila* PCGF3 homolog, L(s)37Ah, interacts with dRING and is required for the majority of H2A118 ubiquitination (58). The single *PCGF* gene in *C*. *elegans* also falls into the PCGF3 family, albeit without high support. It is interesting to note that the PCGF3 family seems to be the group which has been lost less often that any of the other PCGF families, being retained in all genomes that we analyzed other than hydrozoan cnidarians. The reason for this and its relevance to our understanding of PRC1 evolution will only become clear upon further investigation of PCGF3 function in a more diverse set of organisms.

Our finding that anthozoan cnidarians contain the same set of PCGF gene families as vertebrates does not support the hypothesis that the diversification of PRC1 complexes is related to the evolution of vertebrate-specific traits (56, 61). The presence of the different PCGF families in anthozoans provides the opportunity to obtain new insights into the evolution of PRC1 complexes both at the molecular and organismal level. For example, PSc, the *Drosophila* PCGF2/4, has the ability to compact chromatin due to the presence of a repressive C-terminal region. This property can be found in PCGF2/4 proteins from many species, including several invertebrates (80). In vertebrates and plants, however, two unrelated PRC1 subunits, CBX2 and EMF1, respectively, have the same molecular function (11, 80). Thus, it remains ambiguous whether ancestral PCGF proteins had this function or whether it evolved independently in different lineages. Understanding the biochemical activities of PRC1 members from early diverging animal lineages could potentially resolve this. At the organismal level, we see that the *PCGF* genes in *Nematostella* are dynamically and differentially expressed during development. This may indicate that these genes play distinct roles at different developmental stages and/or in different tissues or cell types. It will be interesting in the future to dissect these roles to understand whether the molecular and physiological roles of these genes are conserved in different species. Of the two *PCGF5* paralogs in *Nematostella*, *NvPCGF5a* is highly expressed in the nervous system based on our analysis. In addition, both *Nematostella NvPCGF5a* and *NvPCGF5b* are found to be upregulated in *NvElav1*^+^ neurons at later developmental stages (81). This is striking as, in mammals, *PCGF5* is also highly expressed in the neural progenitors (56, 72) and has been shown to play important functions both during neural differentiation and in the adult nervous system (55, 56). This could suggest an ancestral and conserved function of this gene in the nervous system. A comparably well-developed experimental tool set, including stable transgenics, genome editing and transient knockdown approaches, is available for *Nematostella* (67, 82–85), allowing further investigations on the function and interaction partners of the *Nematostella* PCGF5 proteins that may help to unravel potential functional conservation.

Our analysis failed to resolve the placement of PCGF family genes from other early-branching non-bilaterian lineages (ctenophores, sponges, and placozoans), making their evolutionary history unclear. From our analysis we can confidently say that there were at least five PCGF proteins in the last common ancestor of cnidarians and bilaterians. Whether canonical and non-canonical PRC1 complexes evolved at the same evolutionary stage, or whether one evolved earlier than the other, also remains unclear. The presence of a putative *RYBP* and the absence of either *CBX* of *PHC* homologs in choanoflagellates would favor a hypothesis in which non-canonical PRC1 evolved prior to canonical PRC1 (1). Given the divergent sequence and domain composition of the putative choanoflagellate *S. rosetta RYBP*, we consider it important to validate its potential function as a component of a PRC1 complex experimentally before confidently calling it an RYBP.

In conclusion, we have shown that the *PCGF* family expanded early in animal evolution, before the split of bilaterians and cnidarians, and that therefore, the large diversity of PRC1 complexes seen in vertebrates may have arisen early in animal evolution. Given the extensive losses of *PCGF* genes in the major invertebrate model systems, this places anthozoan cnidarians, particularly *Nematostella vectensis*, as the most technically advanced model in which this complexity can be studied outside vertebrates.

## Materials and Methods

### Homology search

To identify homologs of the genes studied here we used tBLASTn searches with the following as query: For *PCGF* genes we used *Drosophila* PSc, for RING1/2 we used dRING, for CBX genes we used *Drosophila* Pc, for PHC we used *Drosophila* Ph, and for RYBP we used human RYBP. In any case where we could not find any homologs we also used sequences from more closely related groups as a query to confirm. For the majority of species we used the NCBI database. For *Mnemiopsis leidyi* we used the NHGRI *Mnemiopsis leidyi* genome portal (http://research.nhgri.nih.gov/mnemiopsis), for *Schmidtea mediterranea* we used the *Schmidtea mediterranea* genome database (http://smedgd.neuro.utah.edu/), for *Capitella teleta* we used the Joint Genome Institute (https://mycocosm.jgi.doe.gov/Capca1/Capca1.home.html) and for *Hydractinia echinata* sequences were obtained by tBlastn into the transcriptome (https://research.nhgri.nih.gov/hydractinia/download/index.cgi?dl=e_trinity). Genes were designated as orthologs using BLASTp searches with both human and *Drosophila* sequences in the NCBI nr database as well as by analyzing domain composition using Pfam. In a few cases the gene models were obviously incomplete (i.e. very short or missing a domain) and in these cases we extracted the genomic region and performed a de novo annotation to extend the gene models using Augustus (http://bioinf.uni-greifswald.de/augustus/submission). We used the nomenclature as follows: If a gene had already been assigned a name then this was used and the species identifier was added in front. If genes were not already named we named them with the protein name, i.e. PCGF, RING or CBX, preceded by the species identifier and followed by a unique letter (a,b etc.).

### Cloning of *Nematostella* PCGF genes

*Nematostella* PCGF genes were identified as above using the JGI genome browser (http://genome.jgi.doe.gov/Nemve1/Nemve1.home.html) and cloned using standard procedure into pCR4 backbones. In the case of *NvPCGF5a* the sequence was obtained from the NvERTx database (68).

### Phylogenetic analysis

A full list of genes used for phylogenetic analysis can be found in Supplementary File S1. For the PCGF phylogenies, the full-length protein-coding sequences were aligned automatically using MUSCLE v3.8.31 (86). Alignment files are available upon request. ProtTest3 (87), which calls PhyML for estimating model parameters (88), was used to select the best-fit model of protein evolution for each alignment. The best-fit model for the Cnidaria plus Bilateria PCGF and RING1/2 alignment (Fig. S2) was VT + I + Γ + F, where ‘VT’ indicates the substitution matrix, ‘I’ specifies a proportion of invariant sites, ‘Γ’ specifies gamma-distributed rates across sites, and ‘F’ specifies the use of empirical amino acid frequencies in the dataset. The best model for the full taxon set PCGF and RING1/2 alignment (Fig. S3) was WAG + I + Γ + F, where ‘WAG’ indicates the substitution matrix. The best model for the full taxon set PCGF only alignment (Fig. S5) was WAG + I + Γ. Maximum likelihood analyses were performed with RAxML v8.2.9 (89). For each phylogeny, we conducted two independent searches each with a total of 100 randomized maximum parsimony starting trees; we then compared the likelihood values among all result trees and chose the best tree from among these. One hundred bootstrapped trees were computed and applied to the best result tree for each analysis. Bayesian analyses were performed with MrBayes3.2.5 × 64 (90) and the same best fit model of protein evolution from ProtTest3 as described above for each set. Two independent five million generation runs of five chains each were run, with trees sampled every 100 generations. The final ‘average standard deviation of split frequencies’ between the two runs for each phylogeny was always less than 0.05. This diagnostic value should approach zero as the two runs converge and an average standard deviation value between 0.01 and 0.05 is considered acceptable for convergence. In each case, a majority rule consensus tree was produced, and posterior probabilities were calculated from this consensus. Trees were rooted in FigTree v1.3.1 [FigTree, a graphical viewer of phylogenetic trees. http://tree.bio.ed.ac.uk/software/figtree/.]. Bayesian posterior probabilities are shown on the Bayesian trees (Fig. 2), S4, S6).

### Identification and annotation of PCGF cluster

The scaffolds containing the PCGF cluster are as follows: *Aiptaisa*, NW_018385238.1; *Acropora*, NW_015441081.1; *Nematostella*, NW_001834266. The genes upstream and downstream of the *PCGF* genes were identified based on their closest human orthologs based on reciprocal BLASTp searches in the NCBI nr database. We then used a reciprocal blast approach between the species to confirm that in each case the genes in the cluster shown to be the same do in fact represent each other’s closest orthologs in the other species.

### *Nematostella* maintenance

*Nematostella* were maintained at 18-19°C in 1/3 filtered sea water (NM) and spawned as described previously (91). Fertilized eggs were removed from their jelly packages by incubating in 3% cysteine in NM for 20 minutes followed by extensive washing in NM. Embryos were reared at 21°C and were fixed at 16 hours (blastula), 20 hours (gastrula), 30 hours (late gastrula), 48 hours (early planula) and 72 hours (late planula).

### Generation of the *NvPCGF5a*::eGFP transgenic reporter

We amplified ~ 5.3 kb upstream of the *NvPCGF5* coding sequence including the first two introns and 138bp of coding sequence using primers : CACCCCGCAACATGAAGACAAATTG; Rv, TCGGCAAACTAAAAAAAATATATATATATAAATAAG and cloned it in frame with a codon optimized eGFP followed by an SV40 terminator sequence in a pUC57 backbone as previously used (81) using NEB HiFi Mastermix (NEB, EN2621s). Transgenic animals were generated using meganuclease mediated transgenesis as previously described (82).

### Fixation, *in-situ* hybridization and immunofluorescence

Animals were fixed in ice cold 0.2% glutaraldehyde/3.7% formaldehyde in NM for 1.5 minutes followed by 1 hour at 4°C in 3.7% formaldehyde in PBT (PBS + 0.1% tween). Animals were washed several times in PBT and those used for in-situ hybridization were dehydrated through a series of methanol washes and stored in 100% methanol at −20°C. In situ hybridization and immunofluorescence were performed as previously described (75) with the replacement of the DAPI incubation with a 1 hour incubation in Hoechst 33342 (Thermo Fisher Scientific, 62249) at 1:100 for >1 hour. Antibodies used were: anti-dsRed (Clontech, 632496) 1:200, mouse anti-mCherry (Clontech, 632543) 1:200, mouse anti-GFP (Abcam, Ab1218) 1:100, rabbit anti-GFP (Abcam, Ab290) 1:100, goat anti-rabbit Alexa 488 (Life Technologies, A11008), goat anti-rabbit Alexa 568 (Life Technologies, A11011), goat anti-mouse Alexa 488 (Life Technologies, A11001) and goat anti-mouse Alexa 568 (Life Technologies, A11004) 1:200. Samples were imaged on either a Nikon Eclipse E800 compound microscope with a Nikon Digital Sight DSU3 camera or on a Leica SP5 confocal microscope.

## Acknowledgments

We thank members of the Rentzsch lab for discussions and support and Océane Tournière for critical reading of the manuscript. Research in FRs lab was funded by a grant from the University of Bergen and Research Council of Norway (251185/F20) and the Sars Centre core budget. CES was supported by University of Florida Start-up funding.

