## Supplementary material for "The genetic basis for PRC1 complex diversity emerged early in animal evolution": Supplentary figures

File S1. Excel table containing information on all sequences used in the phylogenetic analysis.

Table S1. Polycomb gene names and abbreviations in *Drosophila* and vertebrates.

| ***Drosophila*** | **Vertebrate** |
|---|---|
| **PRC1** | |
| Polycomb(Pc) | Chromobox 2 (CBX2)<br>Chromobox 4 (CBX4)<br>Chromobox 6 (CBX6)<br>Chromobox 7 (CBX7)<br>Chromobox 8 (CBX8) |
| Posterior sex combs(Psc)<br>Suppressor of zeste 2 (Su(z)2) | Polycomb Group RING Finger 1 (PCGF1)<br>Polycomb Group RING Finger 2 (PCGF2)<br>Polycomb Group RING Finger 3 (PCGF3)<br>Polycomb Group RING Finger 4 (PCGF4)<br>Polycomb Group RING Finger 5 (PCGF5)<br>Polycomb Group RING Finger 6 (PCGF6) |
| Polyhomeotic (Ph) | Polyhomeotic Homolog 1 (PHC1)<br>Polyhomeotic Homolog 2 (PHC2)<br>Polyhomeotic Homolog 3 (PHC3) |
| Sex combs extra (Sce or dRING) | RING1 or RING1a<br>RING2 or RING1b |
| **PRC2** | |
| Extra sex combs (Esc) | Embryonic Ectoderm Development (EED) |
| Suppressor of Zeste 12 (Su(z)12) | Polycomb protein SUZ12 (SUZ12) |
| Enhancer of Zeste (E(z)) | Enhancer Of Zeste Homolog 1 (EZH1)<br>Enhancer Of Zeste Homolog 2 (EZH2) |

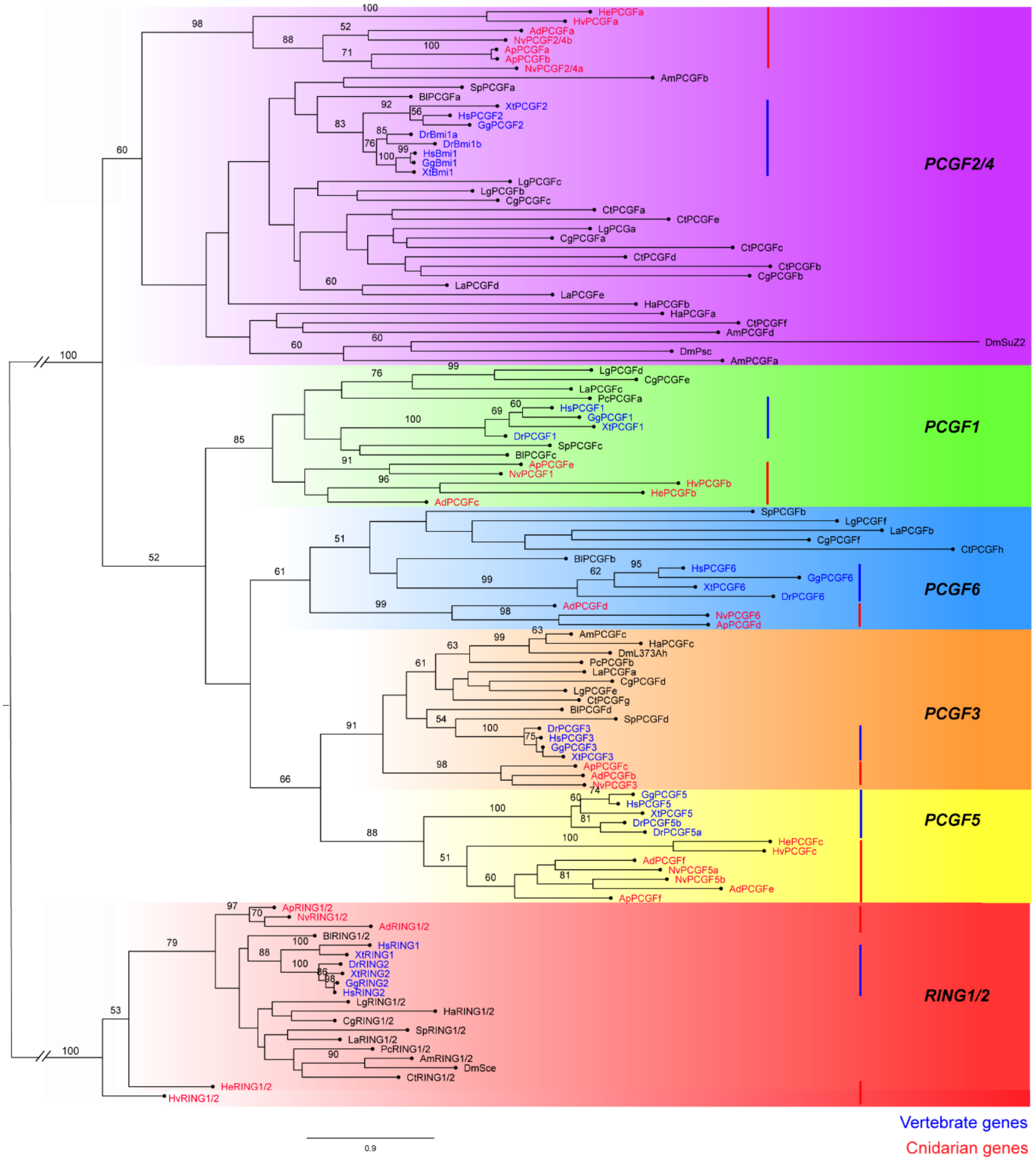


Fig. S1. Phylogenetic tree of cnidarian and bilaterian *PCGF* and *RING1/2* genes according to the maximum likelihood analysis. *RING1/2* genes were used as the outgroup. There were five major families of *PCGF* genes (*PCGF1, 2/4, 3, 5, 6*), which are highlighted by different colored boxes. Numbers above branches correspond to ML bootstrap values. Only values ≥ 50 are shown. Red bars and red font indicate the position of vertebrate genes and blue bars and blue font indicates the position of cnidarian genes. Species names are abbreviated as in Fig. 2.

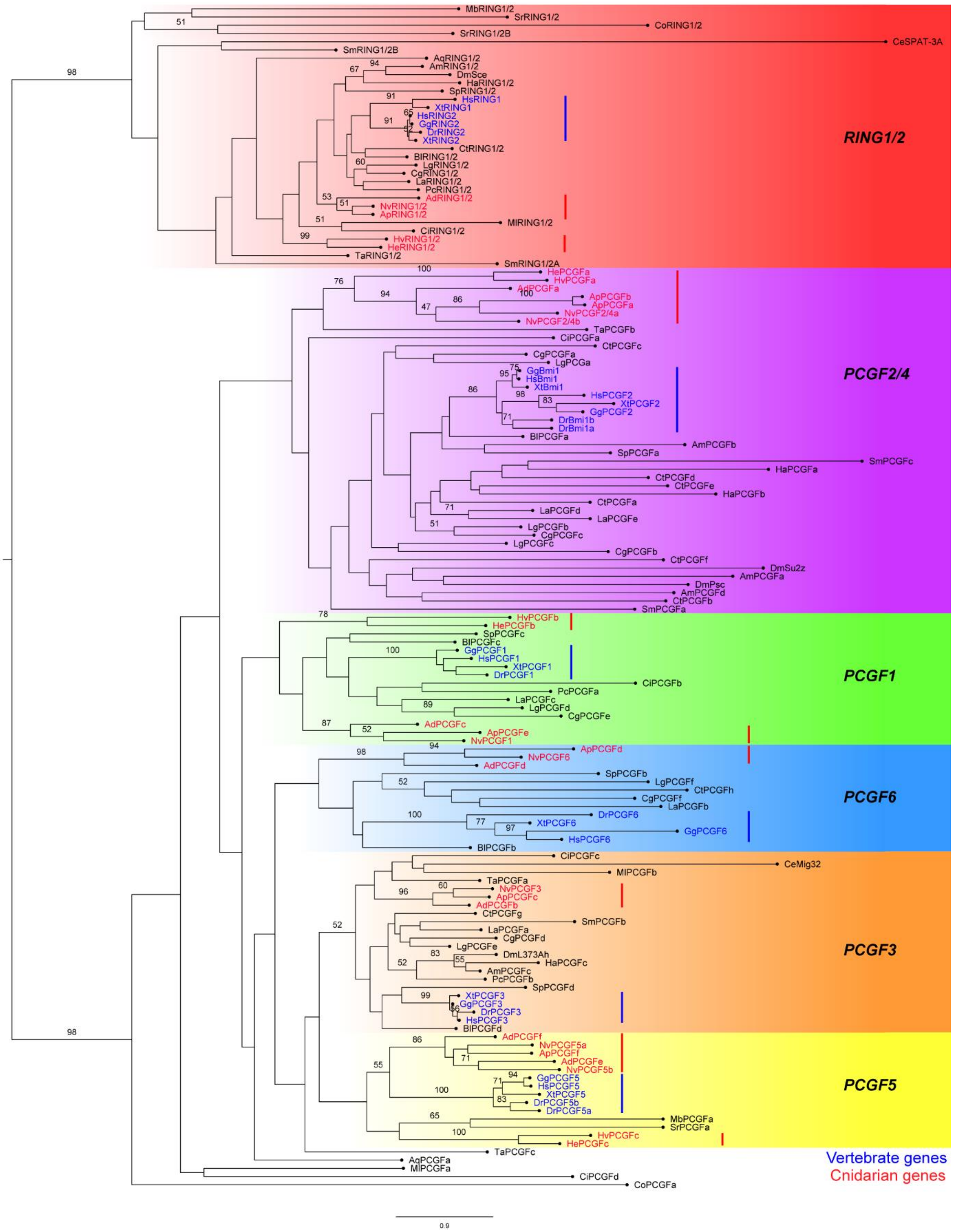


Fig. S2. Phylogenetic tree of the full taxon set of PCGF and *RING1/2* genes according to the maximum likelihood analysis. *RING1/2* genes were used as the outgroup. There were five major families of *PCGF* genes (*PCGF1, 2/4, 3, 5, 6*), which are highlighted by different colored boxes. Five genes were not placed into either RING1/2 or one of the five PCGF families and branched outside our named groups but fall within the overall PCGF group. Numbers above branches correspond to ML bootstraps. Only values ≥ 50 are shown. Red bars and red font indicate the position of vertebrate genes and blue bars and blue font indicates the position of cnidarian genes. Species names are abbreviated as follows: Ad*, Acropora digitifera*; Ap*, Aiptasia pallida*; Aq, *Amphimedon queenslandica*; Am*, Apis mellifera*; Bl*, Branchiostoma floridae*; Ce*, Caenorhabditis elegans*; Cg*, Crassostrea gigas*; Ct*, Capitella teleta*; Co*, Capsaspora owczarzaki*; Ci*, Ciona intestinalis*; Dr*, Danio rerio*; Dm*, Drosophila melanogaster*; Gg, *Gallus gallus*; Ha*, Hyalella azteca*; Hs*, Homo Sapiens*; He*, Hydractinia echinata*; Hv*, Hydra vulgaris*; La*, Lingula anatina*; Lg*, Lottia gigantea*; Ml*, Mnemiopsis leidyi*; Nv*, Nematostella vectensis*; Mb*, Monosiga brevicollis*; Pc*, Priapulus caudatus*; Sr*, Salpingoeca rosetta*; Sm*, Schmidtea mediterranea*; Sp*, Strongylocentrotus purpuratus*; Ta*, Trichoplax adhaerens*; Xt*, Xenopus tropicalis.*

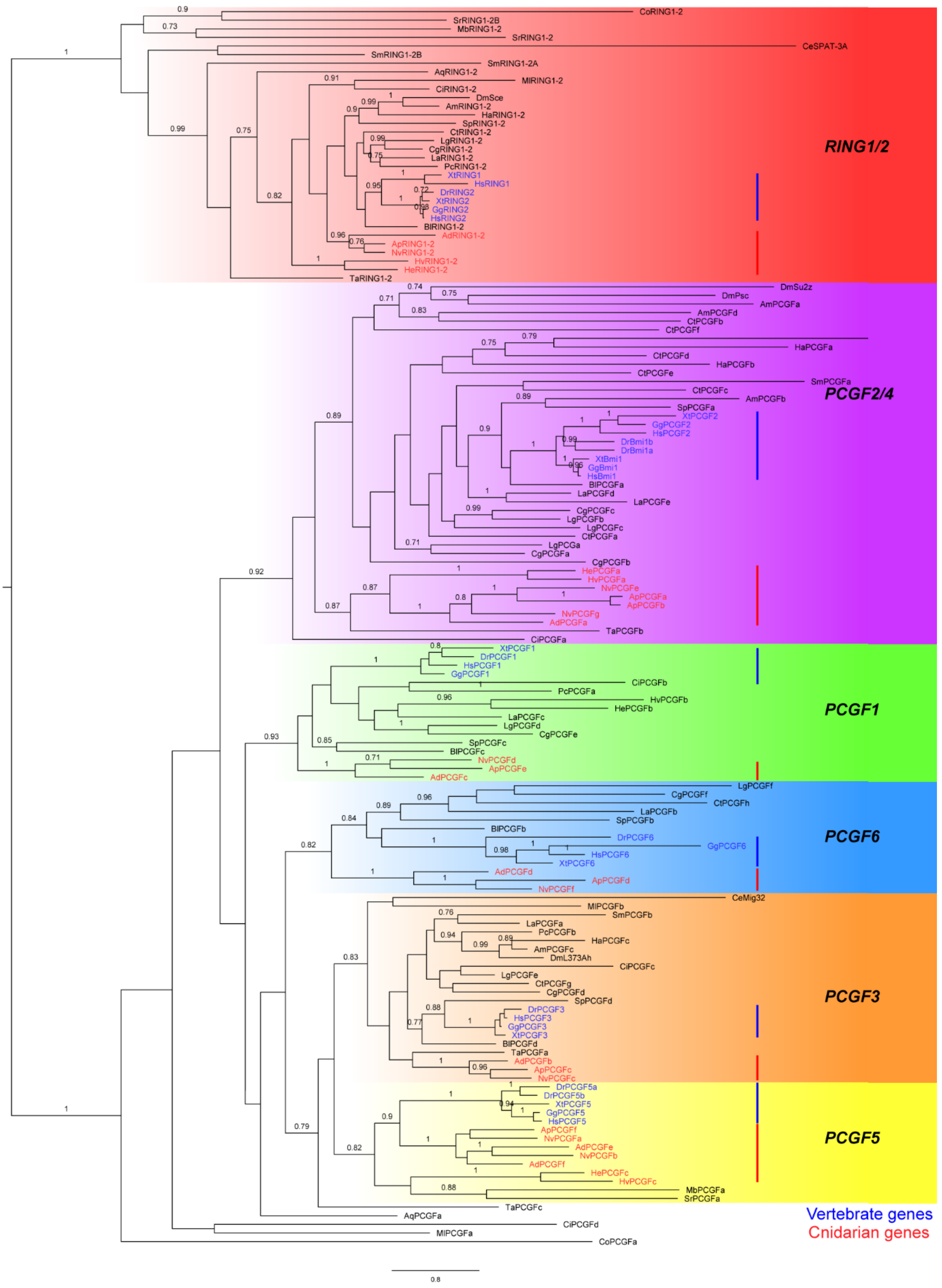


Fig. S3. Phylogenetic tree of the full taxon set of PCGF and *RING1/2* genes according to the Bayesian analysis. *RING1/2* genes were used as the outgroup. There were five major families of *PCGF* genes (*PCGF1, 2/4, 3, 5, 6*), which are highlighted by different colored boxes. Numbers above branches correspond to Bayesian posterior probabilities. Only values ≥ 0.7 are shown. Red bars and red font indicate the position of vertebrate genes and blue bars and blue font indicates the position of cnidarian genes. Species names are abbreviated as in Fig. S2.

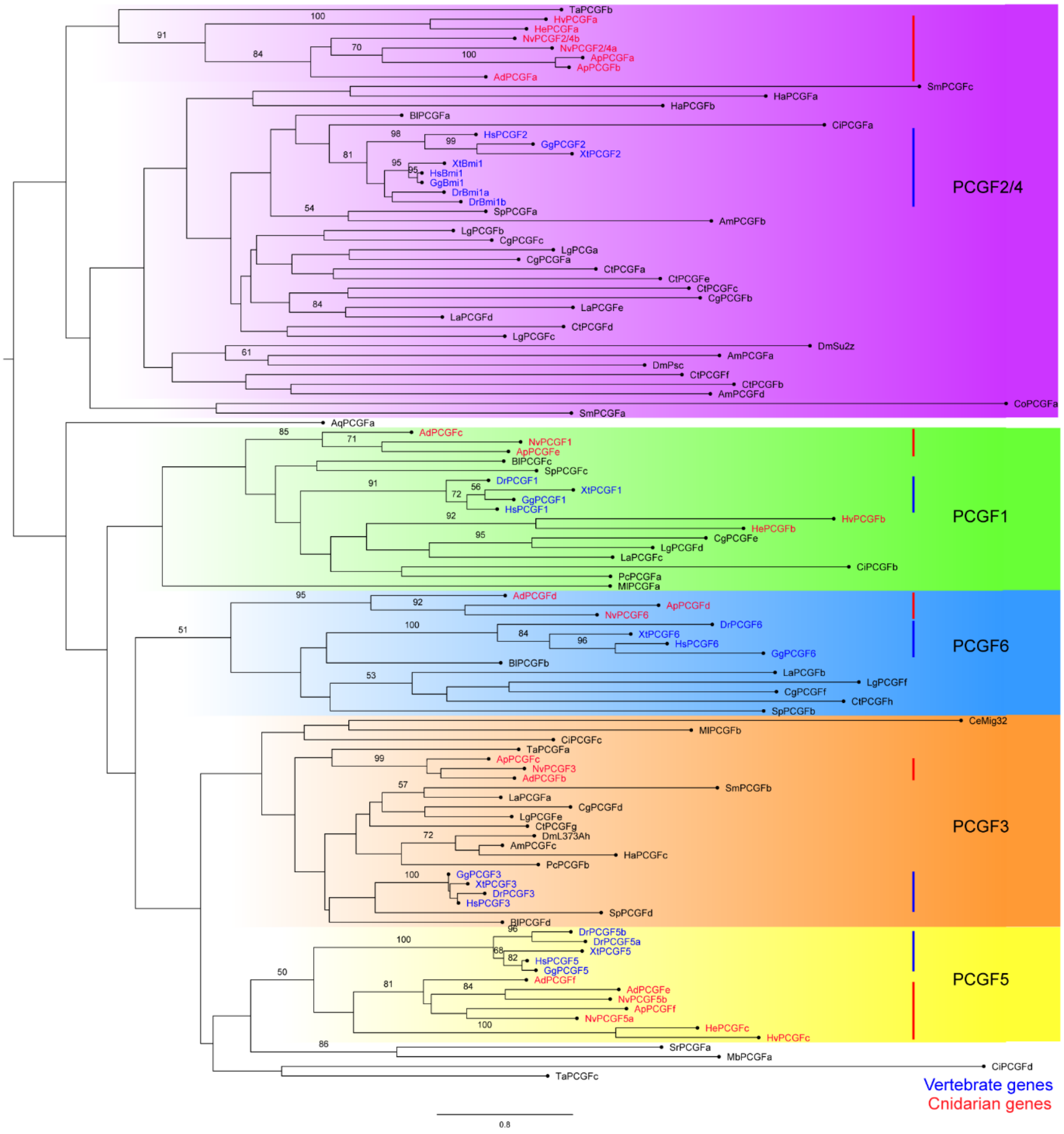


Fig. S4. Phylogenetic tree of the full taxon set of PCGF genes alone according to the maximum likelihood analysis. The tree is midpoint rooted. There were five major families of *PCGF* genes (*PCGF1, 2/4, 3, 5, 6*), which are highlighted by different colored boxes. Numbers above branches correspond to ML bootstraps. Only values ≥ 50 are shown. Red bars and red font indicate the position of vertebrate genes and blue bars and blue font indicates the position of cnidarian genes. Species names are abbreviated as in Fig. S2.

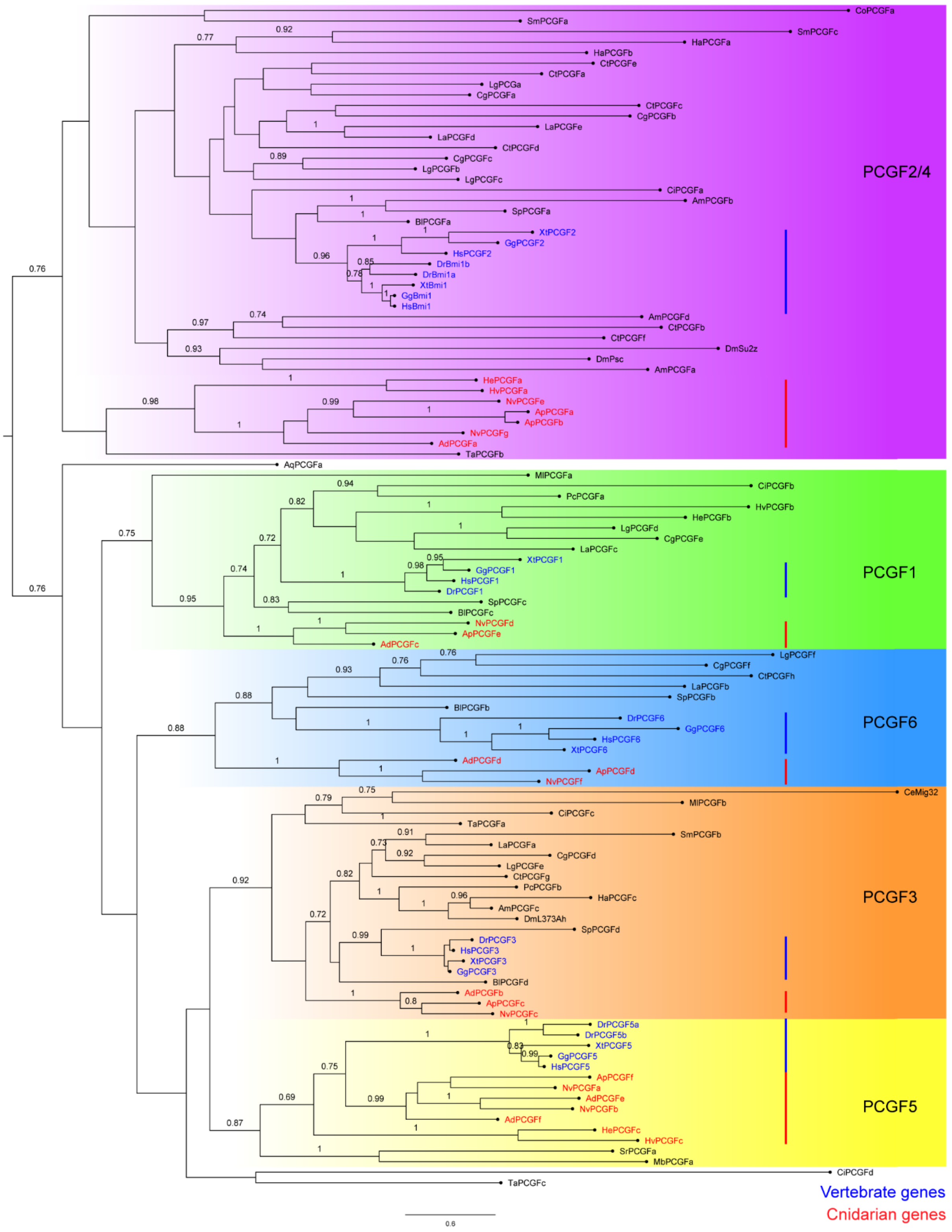


Fig. S5. Phylogenetic tree of the full taxon set of PCGF genes alone according to the Bayesian analysis. The tree is midpoint rooted. There were five major families of *PCGF* genes (*PCGF1, 2/4, 3, 5, 6*), which are highlighted by different colored boxes. Numbers above branches correspond to Bayesian posterior probabilities. Only values

≥ 0.7 are shown. Red bars and red font indicate the position of vertebrate genes and blue bars and blue font indicates the position of cnidarian genes. Species names are abbreviated as in Fig. S2.

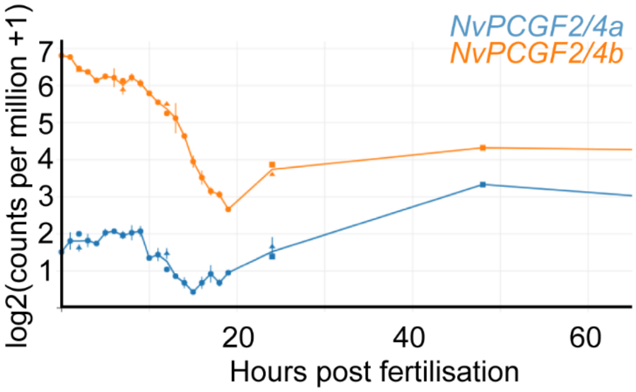


Fig. S6. Expression of canonical *PCGF* genes in *Nematostella vectensis*. (A) Expression analysis of the *Nematostella* canonical *PCGF* genes throughout embryonic development taken from NvERTx database (68).
